# The *Magnaporthe oryzae* MAP kinase Pmk1 regulates polycomb repressive complex 2 to reprogram genes expression for biotrophic growth

**DOI:** 10.1101/2021.04.20.440724

**Authors:** Xuan Cai, Bozeng Tang, Ahmed Hendy, Zhiyong Ren, Caiyun Liu, Muhammad Kamran, Junjie Xing, Lu Zheng, Hao Liu, Junbing Huang, Xiao-Lin Chen

## Abstract

Biotrophic and hemibiotrophic fungi have evolved the ability to colonize living plant cells, but how they establish biotrophic growth by remodeling gene expression is poorly understood. By using *in planta* invasive hyphae (IH) of *Magnaporthe oryzae* to perform an integrated Chromatin immunoprecipitation sequencing (ChIPseq) and RNA-seq analysis, combining with biological and cellular analyses, we found Polycomb repressive complex 2 (PRC2)-mediated epigenetic repression plays a key role in regulating biotrophic growth. ChIPseq for biotrophic IH samples identified 1701 PRC2 target genes. RNA-seq analysis showed that expression of 462 PRC2 target genes were up-regulated in the Δ*suz12* mutant, while 82 were down-regulated, indicating a major role of PRC2 in gene repression of IH. During biotrophic growth, PRC2 repressed fungal cell wall synthesis genes and extracellular enzyme genes required for penetration, and secondary metabolites biosynthesis genes required for necrotrophic growth. A great number of effector-encoding genes were repressed by PRC2, which were highly expressed during penetration stage, suggesting PRC2 coordinates biotrophic growth by regulating effector suppression for immune evasion. This regulation was finely coordinated by Pmk1, through regulating phosphorylation, nuclear localization and protein abundance of Suz12. Our results indicate that the Pmk1-PRC2 regulatory module is required for gene remodeling to facilitate biotrophic growth in *M. oryzae*.

**IMPORTANCE:** Biotrophic and hemibiotrophic fungi establish a biotrophic stage for infection in host cells. For example, *M. oryzae* forms appressoria to penetrate host cell and establish a biotrophic growth stage for infection. How gene expression patterns are elaborately controlled for fungal biotrophic growth is largely unknown. In this study, we found that, the PRC2-mediated H3K27me3 repressed fungal penetration-required cell wall synthesis genes and extracellular enzyme genes, and necrotrophic growth-required secondary metabolites biosynthesis genes for biotrophic growth. Interestingly, a great number of effector-encoding genes were also repressed by PRC2 at biotrophic stage, which were highly expressed at penetration stage, suggesting PRC2 coordinates biotrophic growth by regulating effector suppression for immune evasion. The PRC2-mediated epigenetic repression is therefore required for the gene expression remodeling during fungal infection. This regulation was finely coordinated by Pmk1, through regulating nuclear localization and protein abundance of the PRC2 component Suz12.

## INTRODUCTION

For transcriptional regulation, epigenetic layer histone lysine methylation plays an important role, especially histone H3 Lys 4 methylation (H3K4me)-mediated epigenetic activation and histone H3 Lys 27 methylation (H3K27me)-mediated epigenetic repression (1,2). H3K27me3 is a genetic mark for transcriptional repression, which is commonly modulated by Polycomb group (PcG) proteins. PcG proteins were first genetically identified in *Drosophila*, and required for maintaining the silenced state of developmental regulators of developmental related Hox genes (3). Recent advances have revealed the molecular mechanisms and biological effects of PcG proteins-mediated silencing in different organisms (4,5,6). Using genome-wide chromatin immunoprecipitation (ChIP) analyses, hundreds of PcG target genes were identified, such as in *D. melanogaster* (4,7) and mammalian cells (5,6,8,). Through suppressing different target genes, PcG proteins mediated global silencing system plays key roles in development, stem cell biology and cancer (9,10).

Two main PcG protein complexes, PRC1 (polycomb repressive complex 1) and PRC2 (polycomb repressive complex 2), were found in the previous studies. PRC2 methylates histone H3 on Lys27 (H3K27) to facilitate a feature of chromatin silence (11,12). PRC1 functions as an executor for silencing of target genes (13). In *Drosophila*, PRC2 is composed by several proteins, including catalytic SET domain containing lysine methyltransferase EZH (enhancer of zeste), WD40 domain protein EED (enhanced ectoderm development), suppressor of zeste-12 (Su(Z)12), and nucleosome remodeling factor 55 (NURF55) (14). Components biological functions of PcG, as well as PcG-mediated chromatin mechanisms have been widely studied in stem cell biology, cancer biology and multicellular development in plants and animals (9,15).

However, function studies of PcG in filamentous fungi are relatively less at the present. PRC2 have been identified and studied in non-pathogenic fungus *Neurospora crassa* (16), the plant fungal pathogens *Fusarium graminearum, Fusarium fujikuroi* and *Zymoseptoria tritici* (17,18,19), the human fungal pathogen *Cryptococcus neoformans* (20), and the endophytic symbiont fungus *Epichloё festucae* (21). These studies suggested that PRC2 plays key roles in suppressing of the secondary metabolite metabolism genes.

The filamentous fungus *Magnaporthe oryzae* can infect rice and barley, which has become a model plant fungal pathogen for study of plant-fungus interaction (22). During infection, *M. oryzae* forms appressoria to penetrate cuticle layer of the host plant. After penetration, the penetration peg differentiates to primary hypha, which subsequently differentiates to bulbous secondary hypha and establish a biotrophic stage (23). The fungus then infects neighboring host cells and finally switch into a necrotrophic growth stage. During the early stages of host cell infection, *M. oryzae* secrete effector proteins to suppress host defenses (24,25), but how these effectors are elaborately regulated in expression has not been well determined. How biotrophic growth of *M. oryzae* established is also largely unknown.

Recently, it is reported that *M. oryzae* PRC2 can regulate transcriptional dynamics of effector proteins through transcriptional repression, using *ex planta* experiments (26). In this study, we used *in planta* invasive hyphae (IH) samples to perform a chromatin immunoprecipitation sequencing (ChIP-seq), and show that PRC2-mediated H3K27me3 reprograms transcription of many genes for biotrophic growth of *M. oryzae*. The PRC2-repressed genes are tightly associated with fungal cell wall remodeling, cell wall degrading enzymes (CWDE), and secondary metabolism, which are important for invasive growth after penetration into plant tissue. A great number of effectors expression are also repressed by PRC2, which shows a strong transcription shift between early and late stage during infection. More importantly, we proved the PRC2 is exactly regulated by the MAP kinase Pmk1, which finely revealed Pmk1-downstream regulatory mechanism and PRC2 upstream regulatory mechanism. Together with the previous study by Zhang et al., (26), our study provides strong evidence that PRC2-mediated H3K27me3 is critical for modulating gene expression for biotrophic growth in *M. oryzae*.

## RESULT

### Identification of *M. oryzae* PRC2 complex components

To investigate the genes associated with PRC2 in *M. oryzae*, we firstly determined components of the PRC2 by whole protein BLAST analyses at Ensembl Fungi (http://fungi.ensembl.org/Magnaporthe_oryzae/Info/Index) and identified *M. oryzae* orthologues of the proteins that are involved in PRC2 in *Drosophila melanogaster*. Orthologues of all the components in this pathway can be identified, all of which are single copies in *M. oryzae* (Fig. S1). Many filamentous ascomycetes contain a full complement of PRC2 components, with one homologue each for Ezh, Eed, Suz12 and the ubiquitous Nurf-55/RbAP46/48 (27). Here, we focus on *M. oryzae* PRC2 complex genes, *MoEED1*, *MoEZH2* and *MoSUZ12*. Domain analysis showed that MoEed1 protein contains four WD40 domains. MoEzh2 protein contains the SET motif, which is important for substrate recognition and enzymatic activity. MoSuz12 contains a zinc-finger C2H2 domain, VEFS box domain and a PHD finger domain (Fig. S2). To determine function of PRC2 in *M. oryzae*, the gene replacement approach was used to generate the null mutants of *MoEED1*, *MoEZH2* and *MoSUZ12* (Fig. S3).

### PRC2 complex is required for vegetative growth and conidiation

To investigate the role of PRC2 complex in the vegetative growth in *M. oryzae*, the colony morphology of the Δ*eed1*, Δ*ezh2* and Δ*suz12* mutants on oatmeal tomato agar plate (OTA) was observed. The colony sizes of the Δ*eed1*, Δ*ezh2* and Δ*suz12* mutants were similar and noticeably reduced compared to the wild-type (Fig. S4A and B). The hyphal tip morphology of the PRC2 complex mutants had been stained with Calcofluor White (CFW), and we found that the average length of apical hyphal cells was significantly reduced compared to that of the wild-type strain (Fig. S4C and D). The conidiation capacity of the Δ*eed1*, Δ*ezh2* and Δ*suz12* mutants were around 44% less than that of the wild-type strain (Fig. S4E and F).

### PRC2 complex is involved in virulence and biotrophic growth

To determine whether deletion of the PRC2 complex affects the infection capacity, we examined the virulence of the wild-type, the PRC2 complex mutant strains on susceptible rice seedlings (*Oryzae sativa* cv. LTH). Conidial suspensions (1 × 10^5^ conidia/mL) of the above strains were respectively sprayed onto rice seedlings. The PRC2 complex mutant genes showed a dramatical reduction of lesion size and number compared to that of the wild-type strain (Fig. 1A). For more evidence, one-week-old barley leaves (*Hordeum vulgare* cv. E9) were also inoculated by spraying a conidial suspension of these strains and a similar result was observed. We also inoculated the mycelial agar plugs onto the wounded rice leaves, which were scratched with a needle. Then we found that lesions caused by the PRC2 complex mutant genes spread less than that produced by the wild-type strain (Fig. 1B), indicating that invasive growth of the PRC2 complex mutants was evidently blocked. We conclude that the PRC2 complex are important regulator of virulence during *M. oryzae* infection.

**FIG 1.**
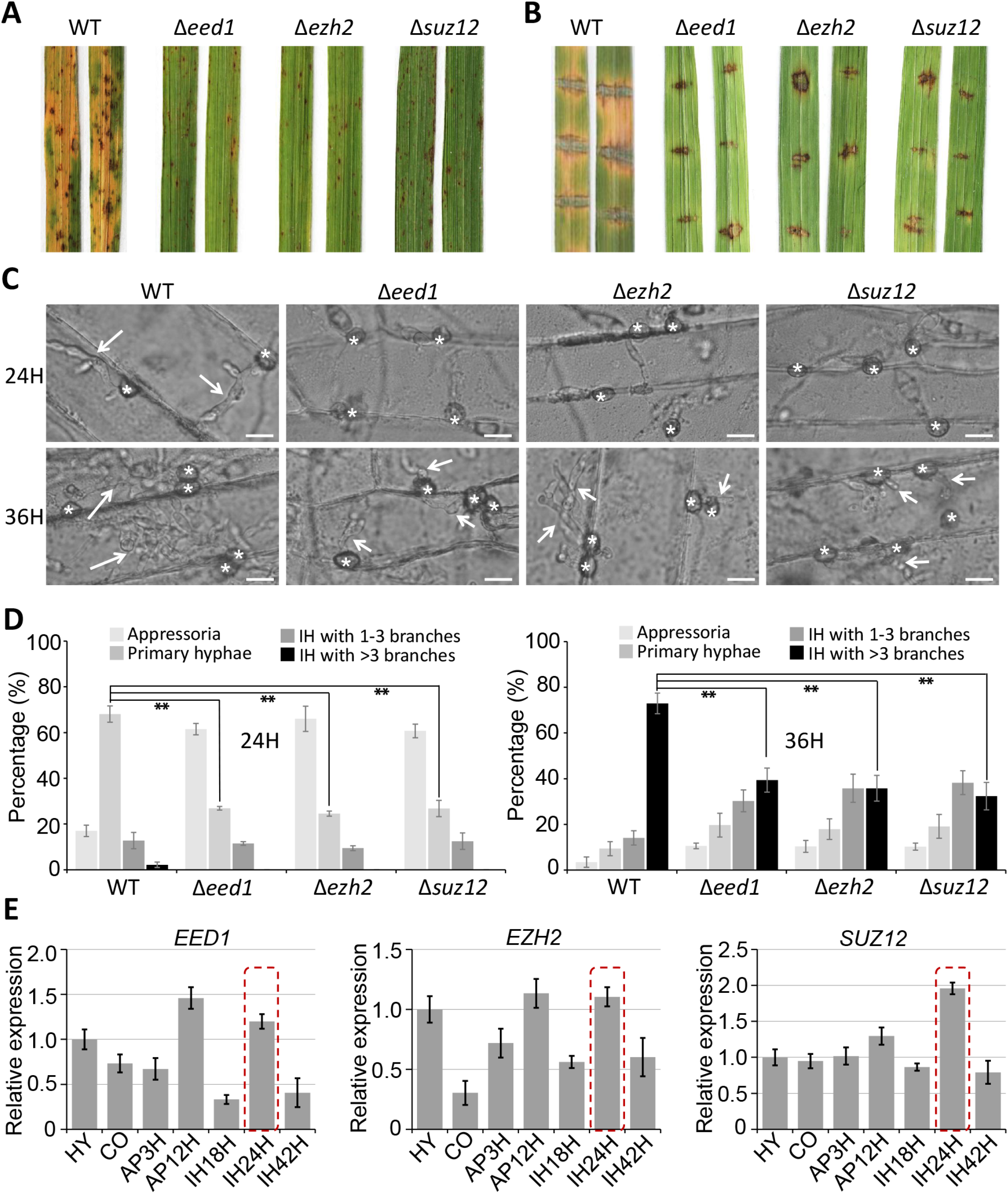
Deletion mutants of the PRC2 complex genes significantly reduced in virulence. (A) Virulence test on rice seedlings. Rice (*Oryza sativa* cv. LTH) leaves were sprayed with conidia suspensions (1 × 10^5^ conidia/ml) and typical leaves were photographed at 5 dpi. (B) Virulence test on wounded rice. Rice leaves were slightly wounded by a needle before inoculating with a mycelial block. Typical leaves were photographed at 3 dpi. (C) Penetration and infectious hyphae were examined at 24 and 36 hpi. Stars indicate the appressoria, and arrows indicate invasive hyphae. AP, PH, and BH indicate appressoria, primary IH, and branched IH, respectively. Bar = 20 mm. (D) Percentages of appressoria with distinct types of IH, primary IH, IH with 1-3 branches, and IH with more than three branches. Means and standard errors were calculated from three independent replicates (n > 100). (E) Expression of PRC2 complex genes at different developmental stages. Relative abundance was normalized by MoTub1. The expression level of each gene in mycelium was set as 1, expressions of other stages were relative to the mycelium stage. Means and standard errors were calculated from three independent replicates (n > 100). MY, mycelium; CO, conidium; AP3H and AP12H, appressorium at 3 hpi and 12 hpi; IH18H, IH24H and IH42H, invasive hyphae at 18 hpi, 24 hpi and 42 hpi.

To further investigating why deletion of the PRC2 complex resulted in the reduction of virulence, we performed live-cell imaging to monitor growth of invasive hyphae. At 24 hpi and 36 hpi, the Δ*eed1*, Δ*ezh2* and Δ*suz12* mutants were severely arrested in biotrophic growth (Fig. 1C and D), indicating the PRC2 complex plays a key role in biotrophic growth in *M. oryzae*.

### PRC2 complex regulates stress response

To determine whether histone tri methylation of H3K27 is involved in stress response, we investigated the sensitivity of the Δ*eed1*, Δ*ezh2* and Δ*suz12* mutants to different kind of stresses. The wild-type and the mutant genes strains were inoculated onto the complete medium (CM) plates supplemented with different reagents and grown for 120 h. The results indicated that the Δ*eed1*, Δ*ezh2* and Δ*suz12* mutants were more sensitive to a series of stresses, especially the cell wall-disturbing reagents (Fig. 2A). Under conditions of 0.1 mg/mL Calcofluor White (CFW), 0.2 mg/mL Congo Red (CR) or 0.005% sodium dodecyl sulphate (SDS), significant reduction of the colony growth happened as a result of the high sensitivity to these cell wall-disturbing reagents. It also found that the sensitivity to other stresses had been increased, including osmotic stress (0.5 M NaCl) and oxidative stress (10 mM H_2_O_2_) (Fig. 2A). These data suggest that PRC2 complex genes are involved in responding to various stresses.

**FIG 2.**
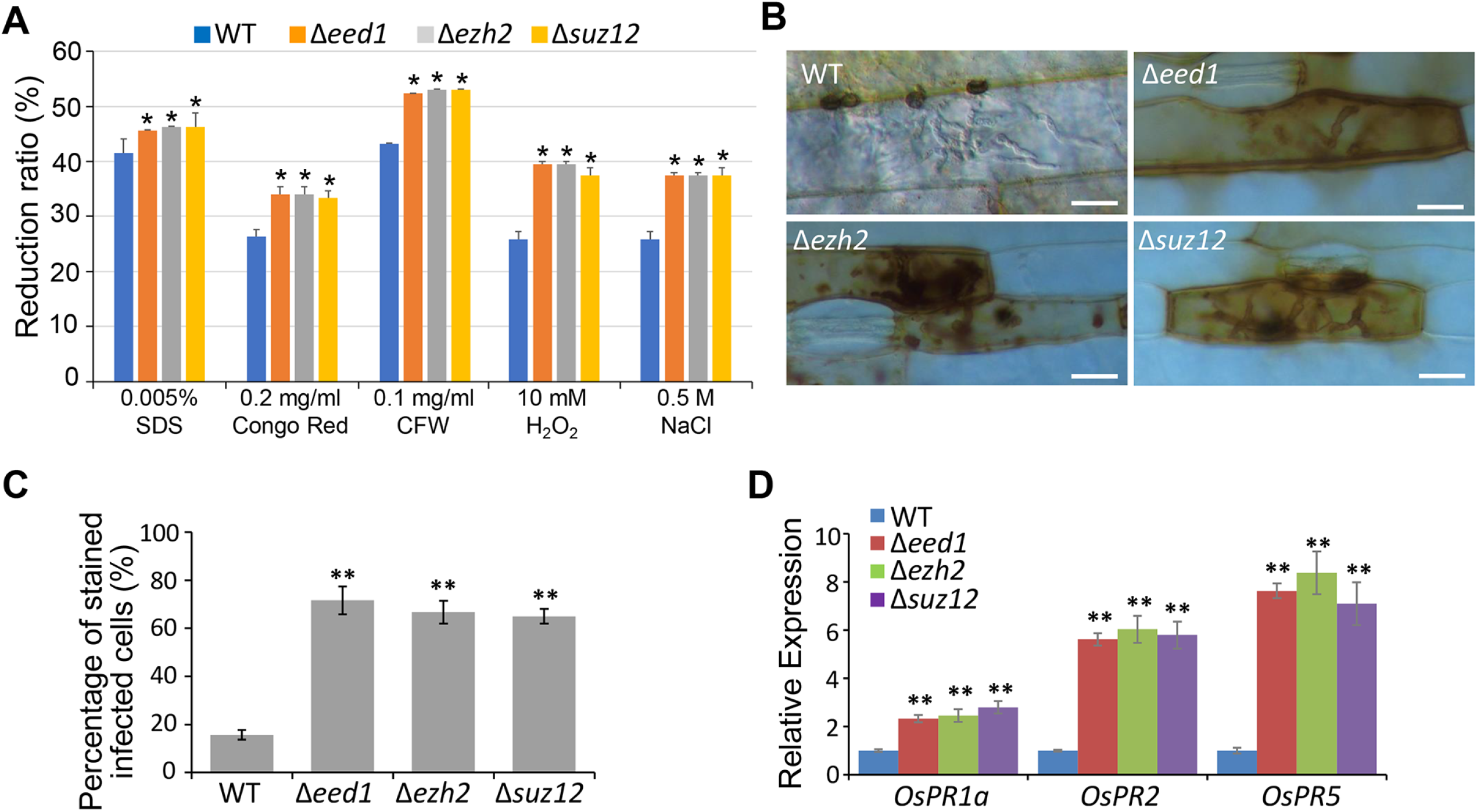
Deletion mutants of PRC2 complex genes are sensitive to different stresses and induces host defense response. (A) Colony growth reduction ratio of PRC2 complex genes’ deletion mutants at different stress conditions. CM plates were supplemented with different stress agents (0.2 mg ml^−1^ Congo Red, 0.1 mg ml^−1^ Calcofluor White (CFW), 0.005% sodium dodecyl sulfate (SDS), 0.5 M NaCl, or 10 mM H_2_O_2_, buffered at pH 8.0). The colonies were photographed at 5 d post inoculation (dpi). Means were calculated from three independent replicates. Significant differences are indicated: *, P < 0.05. (B) DAB staining assay of the penetrated plant cells. Barley epidermis was stained with 3,30-diaminobenzidine at 36 hpi and observed under a microscope. Bars, 20 μm. (C) Percentage of DAB-stained infected cells. Significant differences are indicated: **, P < 0.01. (D) Expression levels of PR genes in rice infected by *M. oryzae* at 36 hpi. Means were calculated from three independent replicates.

### Disruption mutants of PRC2 complex genes induce ROS and host PR gene expression

Invasive growth impairment in *M. oryzae* could be due to a failure to suppress or evade the host immunity. In order to determine whether PRC2 is involved in suppressing or evading host immunity, we subsequently tested whether the Δ*eed1*, Δ*ezh2* and Δ*suz12* mutants could activate host immune responses. Plant immunity responses are often associated with the accumulation of reactive oxygen species (ROS) (28). ROS production was detected with 3,30-diaminobenzidine (DAB) in cells of barley leaves and rice sheaths inoculated with the WT, the Δ*eed1*, Δ*ezh2* and Δ*suz12* mutants. As a result, massive ROS accumulation was commonly detected in barley epidermal cells and rice sheath cells infected by the Δ*eed1*, Δ*ezh2* and Δ*suz12* mutants but not detected in most of the cells infected by the WT strain (Fig. 2B and C). These results suggest that the disruption of PRC2 complex genes could lead to the failure of suppressing host ROS production and thus enhance plant immune responses during infection. To confirm that the disruption of PRC2 complex genes enhances the host immune response, we determined the transcript levels of several infection responsive genes, including OsPR1a, OsPR2 and OsPR5. As expected, expressions of these genes were significantly elevated in the Δ*eed1*, Δ*ezh2* and Δ*suz12* mutant strains, compared with which in the WT (Fig. 2D). Thus, PRC2 may help to evade host innate immunity during fungal infection.

### PRC2 complex is involved in H3 histone tri-methylation (H3K27me3)

The PRC2 complex which contain EED1-SUZ12-EZH2 are required for methylation of H3K27. We therefore wondered if H3K27me3 levels are altered in the Δ*eed1*, Δ*ezh2* and Δ*suz12* mutants. We isolated total histones from wild type and the Δ*eed1*, Δ*ezh2* and Δ*suz12* mutants, and performed Western blots using antibodies of anti-H3K27me1, anti-H3K27me2, and anti-H3K27me3, respectively. We found that the H3K27me3 modification completely disappeared in the Δ*eed1*, Δ*ezh2* and Δ*suz12* mutants comparing with which in the wild-type (Fig. 3A). No difference of H3K27me1 was detected, but H3K27me2seems strengthened in the Δ*eed1*, Δ*ezh2* and Δ*suz12* mutants (Fig. 3A). These results suggest that PRC2 is the major histone methyl transferase responsible for H3K27me3 in *M. oryzae*.

**FIG 3.**
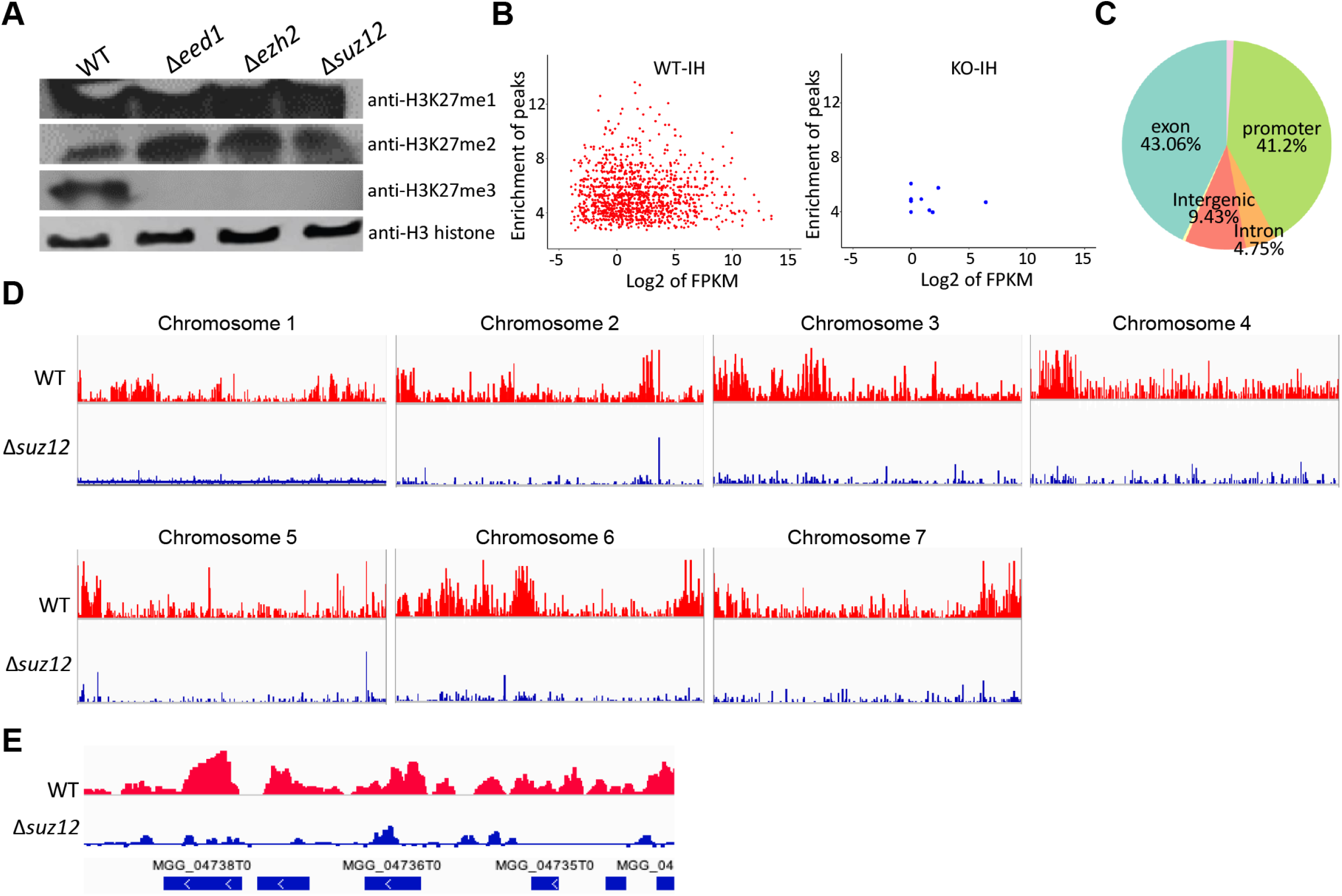
Polycomb complex is involved in histone H3K27 tri-methylation. (A) Western blotting analysis of histone H3K27 modifications in PRC2 gene’s mutants. Total protein extracted from *M. oryzae* cells was subjected to 15% SDS polyacrylamide gel electrophoresis, and probed with antibodies against H3K27me3, and H3 histone. (B) Enrichment peaks of H3K27me3 modification in 24 hpi-invasive hyphae samples of WT (WT-IH) and Δ*suz12* mutant (KO-IH). (C) Distribution of histone modifications across the complete set of *M. oryzae* genes. (D) Over view of ChIP-seq results in *M. oryzae* chromosomes. Blue and red peaks in the wiggle plot represents the normalized ChIP-seq read coverage. (e) Distribution of H3K27me3 enriched peaks in representative genes.

### ChIP-seq analysis of WT and Δ*suz12* using anti-H3K27me3

To further confirm that PRC2 is involved in H3K27me3 modification, we performed ChIP-seq with antibody against histone H3K27me3, using invasive hyphae of the wild type and Δ*suz12* mutant infecting in barley epidermis cells at 24 hpi. We found that the histone H3K27me3 mark cooccurred at most genes using a high-confidence binding threshold. From the ChIP-seq analysis, we identified 1701 positions related to 1272 genes are enriched by histone modification of H3K27me3 in WT, indicating that around 10% of *M. oryzae* 13,000 protein-coding genes are potentially affected by H3K27me3 at biotrophic growth stage (Table S1; Fig. S5). Plotting raw enrichment ratios for genes associated H3K27me3 demonstrates that *SUZ12* binding represents PRC2 binding at almost all target genes (Fig. 3B). Only 11 sites are enriched in Δ*suz12*, suggesting the PRC2 is critical and essential for H3K27me3. We analyzed location of these H3K27me3 peaks in whole genome, and found that most of them were located in gene promoter (43.06%) or exon region (41.2%), while only 4.75% were located in intron region, as well as 9.43% located in intergenic region (Fig. 3C). The sites occupied by H3K27me3 were then mapped throughout the entire genome, which clearly showed that nearly all the H3K27me3 sites in the Δ*suz12* were significantly reduced enrichment (Fig. 3D and E).

### PRC2 complex regulates expression of thousands of genes during biotrophic growth

We examined whether changes in the H3K27me3 were associated with gene activation or silencing during biotrophic growth in *M. oryzae* at a global scale. We performed comparative transcriptomics using same IH samples as in ChIP-seq. After analyzing RNA-seq by DESeq2 package, a total of 2,791 genes showed significant increases (1,493 genes) or decreases (1,298 genes) (FDR < 0.05) in expression levels at 24 hpi IH stage in the Δ*suz12* mutant (Fig. 4A and D; Fig. S6; Table S2). Interestingly, the number of genes up-regulated in the Δ*suz12* mutant was comparable to the number of down-regulated genes, suggesting that PRC2 directly or indirectly plays a role in gene repression, as well as in gene activation for biotrophic growth.

**FIG 4.**
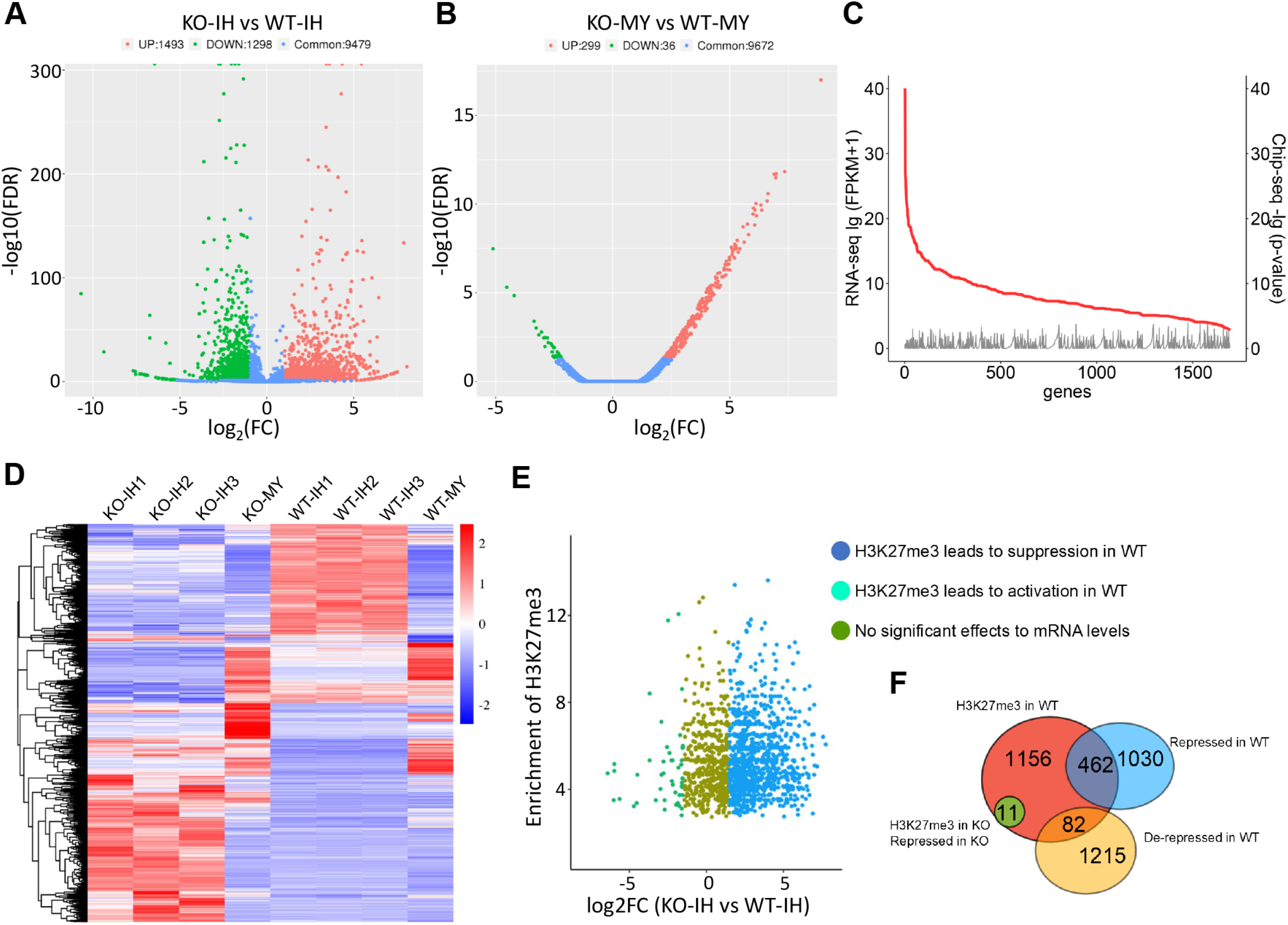
H3K23me3 abundance was generally associated with transcription. (A) Volcano plot for differential gene expression in 24 hpi-invasive hyphae of the Δ*suz12* mutant (KO-IH) comparing with that in WT (WT-IH). (B) Volcano plot for differential gene expression in mycelia of the Δ*suz12* mutant (KO-MY) comparing with that in WT (WT-MY). (C) H3K427me3 abundance was generally negative-associated with gene’s transcription level. Red line indicates the H3K427me3 abundance, and gray line indicates gene expression level. (D) Heatmap shows differential expressed genes via the RNA-Seq FPKM values. KO-IH, 24 hpi-invasive hyphae of the Δ*suz12* mutant; WT-IH: 24 hpi-invasive hyphae of WT; KO-MY, mycelium of the Δ*suz12* mutant; WT-MY: mycelium of WT. (E) Scatter plot shows transcription level changes of the H3K427me3 modified genes. (F) Venn diagram shows overlap of ChIP-seq and RNAseq data identified genes.

However, PRC2-mediated histone H3K27 methylation promotes gene silencing at the majority of its target genes throughout the genome, as demonstrated by correlation between H3K27me3 modification and gene expression (Fig. 4C). We found that in total of 1701 H3K27me3 occupied genes, 462 genes (27.2%) were repressed at biotrophic growth stage, whose expression were significantly increased in the Δ*suz12* mutant. In contrast, only 82 genes (4.82%) were activated at biotrophic growth stage, whose expression were down-regulated in the Δ*suz12* mutant (Fig. 4D and F). These results are consistent with the model that PRC2-mediated histone H3K27 methylation promotes gene silencing at the majority of its target genes. Interestingly, in mycelium samples, only 335 genes showed significant increases (299 genes) or decreases (36 genes) (FDR < 0.05) expression levels in the Δ*suz12* mutant (Fig. 4B and D; Table S2), indicating that PRC2 regulates much less gene’s expression in mycelium compared with biotrophic growth stage.

### Functional classification PRC2-targeted genes

To understand the functions of PRC2 in biotrophic growth stage, we classified the PRC2 target genes into different groups by Gene Ontology (GO) annotation and performed enrichment analysis. GO terms associated with protein secretion, and substance releasing, are statically enriched by PRC2-targeted genes (Fig. 5A). Biological processes and molecular functions analysis both indicated that proteins involved in polysaccharide catabolic process and hemicellulose metabolic process were also enriched, indicating cell wall synthesis and extracellular plant cell wall digestion enzymes were regulated by PRC2 (Fig. 5A). More interestingly, this analysis also indicated that secondary metabolites transport, including antibiotic transport, drug transmembrane transport, and xenobiotic transport, were also regulated by PRC2 (Fig. 5A). To investigate effects of PRC2 to fungal metabolism, further pathway enrichment analysis was performed. This highlighted the regulation of PRC2 in secondary metabolites metabolisms, including the phenylalanine metabolism, aflatoxin biosynthesis, dioxin degradation, polycyclic aromatic hydrocarbon degradation, naphthalene degradation, fluorobenzoate degradation, betalain biosynthesis, and cytochrome P450-mediated metabolisms (Fig. 5B). These analyses indicated that the PRC2 plays important roles in secondary metabolites metabolism.

**FIG 5.**
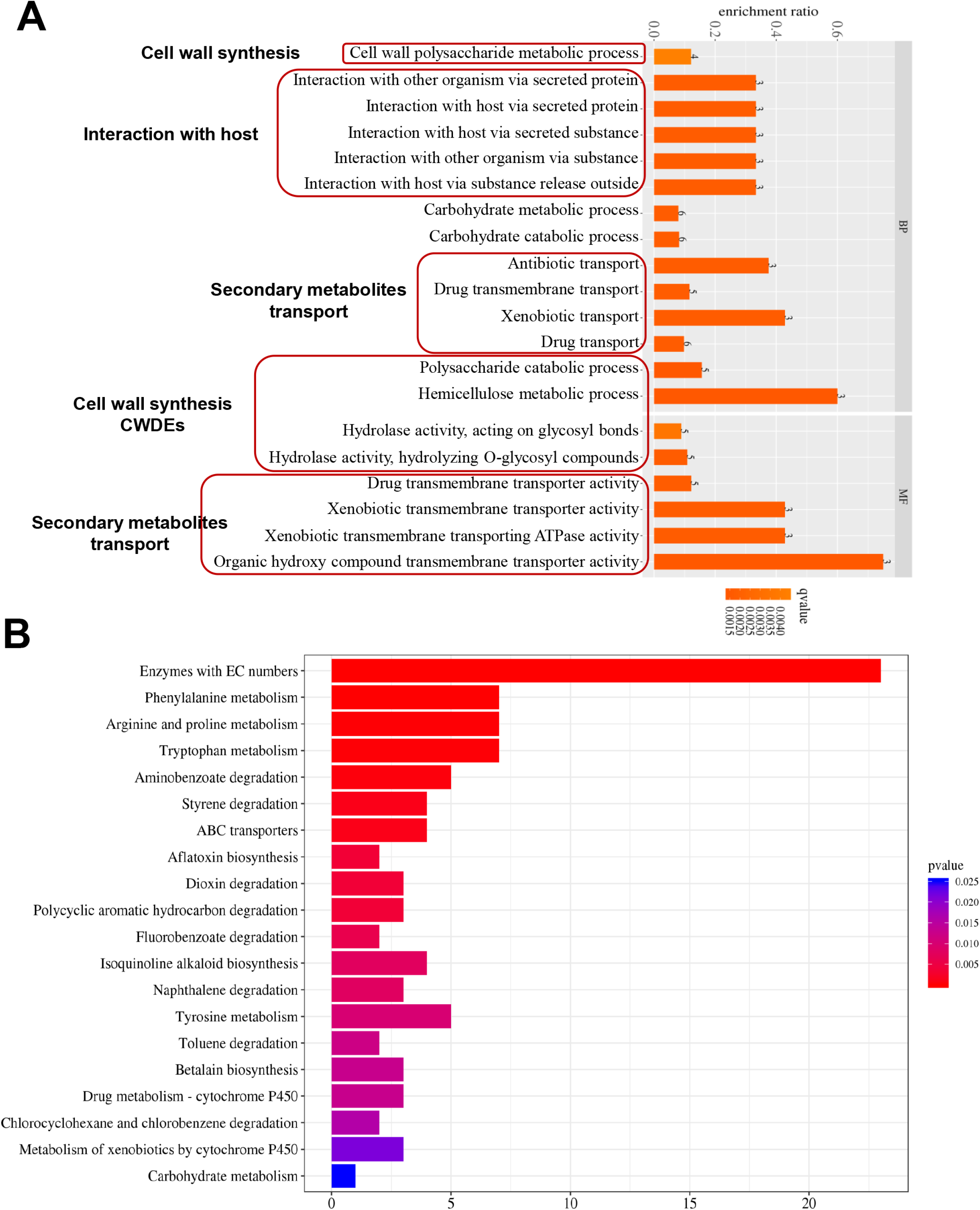
Gene Ontologies and metabolic pathways modulated by H3K27me3. (A) GO enrichment functional classification of the peak-linked genes. (B) KEGG pathway analysis of the peak-linked genes.

### PRC2 coordinates different cellular processes for biotrophic growth

Examination of the targets of PRC2 revealed that they were remarkably enriched for genes of regulators such as transcription factors and protein kinases (Fig. 6A and B; Table S3), which play key roles in regulating downstream cellular and biological processes. Transcription factors, at least four Cys_2_-His_2_ (C2H2) transcription factors, and five Zn_2_Cys_6_ (C6) transcription factors (29,30), were identified as PRC2 targets (Fig. 6A). These included genes for transcription factors known to be important for conidia formation during necrotrophic growth stage (29,30). We found that *ZFP6, GCF8, FZC44, FZC12* and *ZFP11* were severely repressed by PRC2 (Fig. 6A). At least seven protein kinase genes were modified by PRC2-mediated H3K27me3 modification at biotrophic growth stage. Most of these kinase genes were repressed by PRC2 (Fig. 6B), including *MoMCK1*, encoding a key regulator of cell wall biosynthesis and remodeling that is important for host penetration (31). Besides *MoMCK1*, we found that some other important fungal cell wall synthesis genes were also repressed in biotrophic growth stage by PRC2, including genes encoding β-1,3-endoglucanase, alpha-1,2-mannosyltransferase, alpha-1,6-mannosyltransferase, glycosyl hydrolase, glycosyltransferase, and chitin deacetylase (CDA1) and agglutinin isolectin (Fig. 6C; Table S3). All these cell wall-related key genes were significantly reduced expression in the Δ*suz12* mutant, indicating an obvious PRC2-mediated suppression of cell wall integrity during biotrophic growth stage. Cell wall degrading enzyme genes (CWDE) play key roles during fungal penetration process, through digesting plant cell wall (32). Around 20 CWDEs were modified by PRC2-mediated H3K27me3, and at least one half of which were significantly suppressed at biotrophic growth stage (Fig. 6D; Table S3). Collectively, PRC2-mediated H3K27me3 suppresses key regulators of cell wall-related genes and CWDE genes at biotrophic growth stage for successful infection.

**FIG 6.**
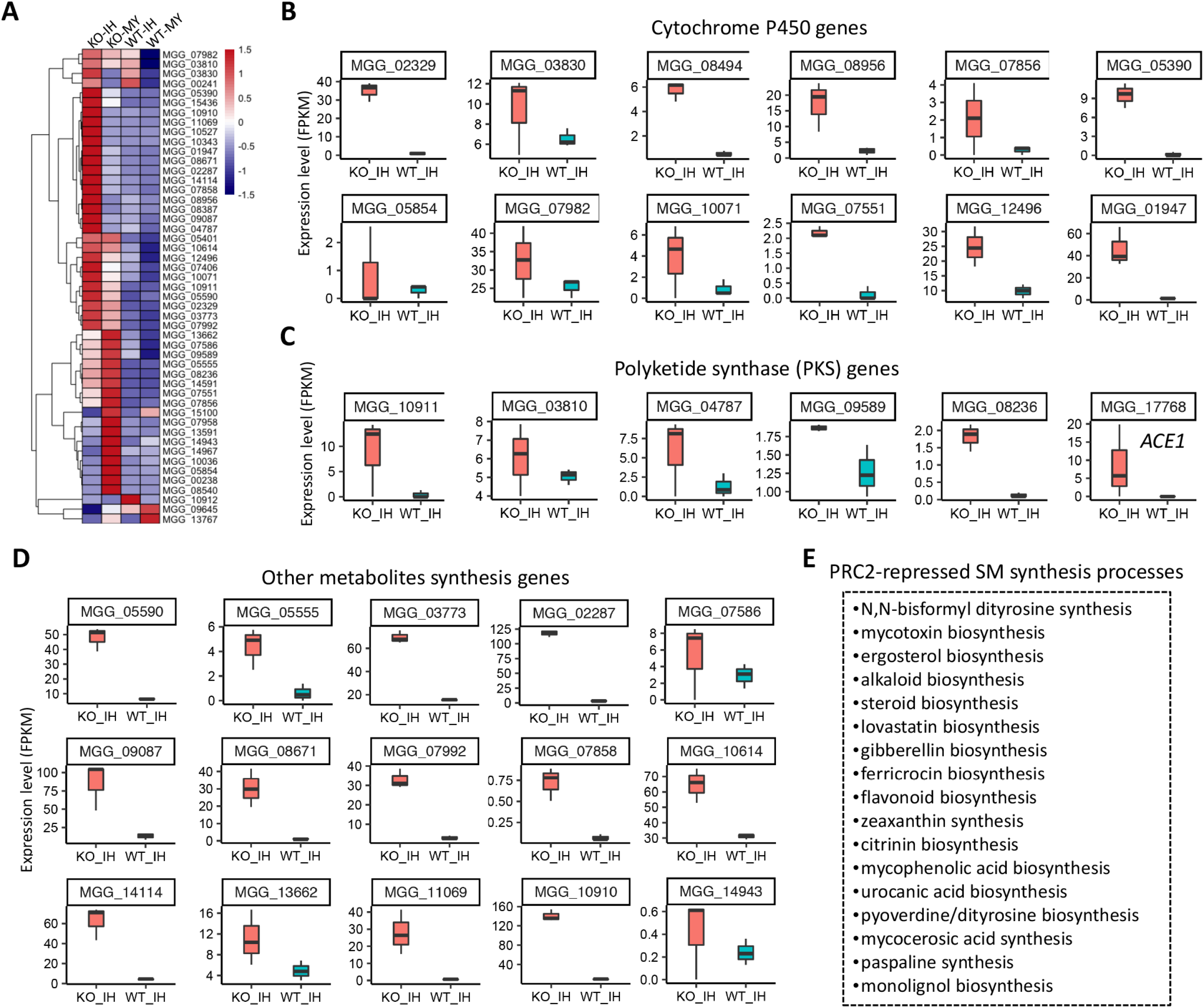
PRC2 represses transcription factor, protein kinase, cell wall synthesis and extracellular enzyme genes. Heatmap shows expression of transcription factor (A), protein kinase (B), cell wall synthesis (C), and extracellular enzyme genes (D). KO-IH, 24 hpi-invasive hyphae of the Δ*suz12* mutant; WT-IH: 24 hpi-invasive hyphae of WT; KO-MY, mycelium of the Δ*suz12* mutant; WT-MY: mycelium of WT.

### PRC2-regulated genes are tightly associated with secondary metabolites biosynthesis

A large number of secondary metabolites biosynthesis pathway genes were identified as PRC2 targets in *M. oryzae* (Table S3). It is obviously that these genes were overall repressed in both 24 hpi invasive hyphae and mycelia samples, but their expressions were significantly increased in the Δ*suz12* mutant either in 24 hpi invasive hyphae sample or in mycelia sample (Fig. 7A).

**FIG 7.**
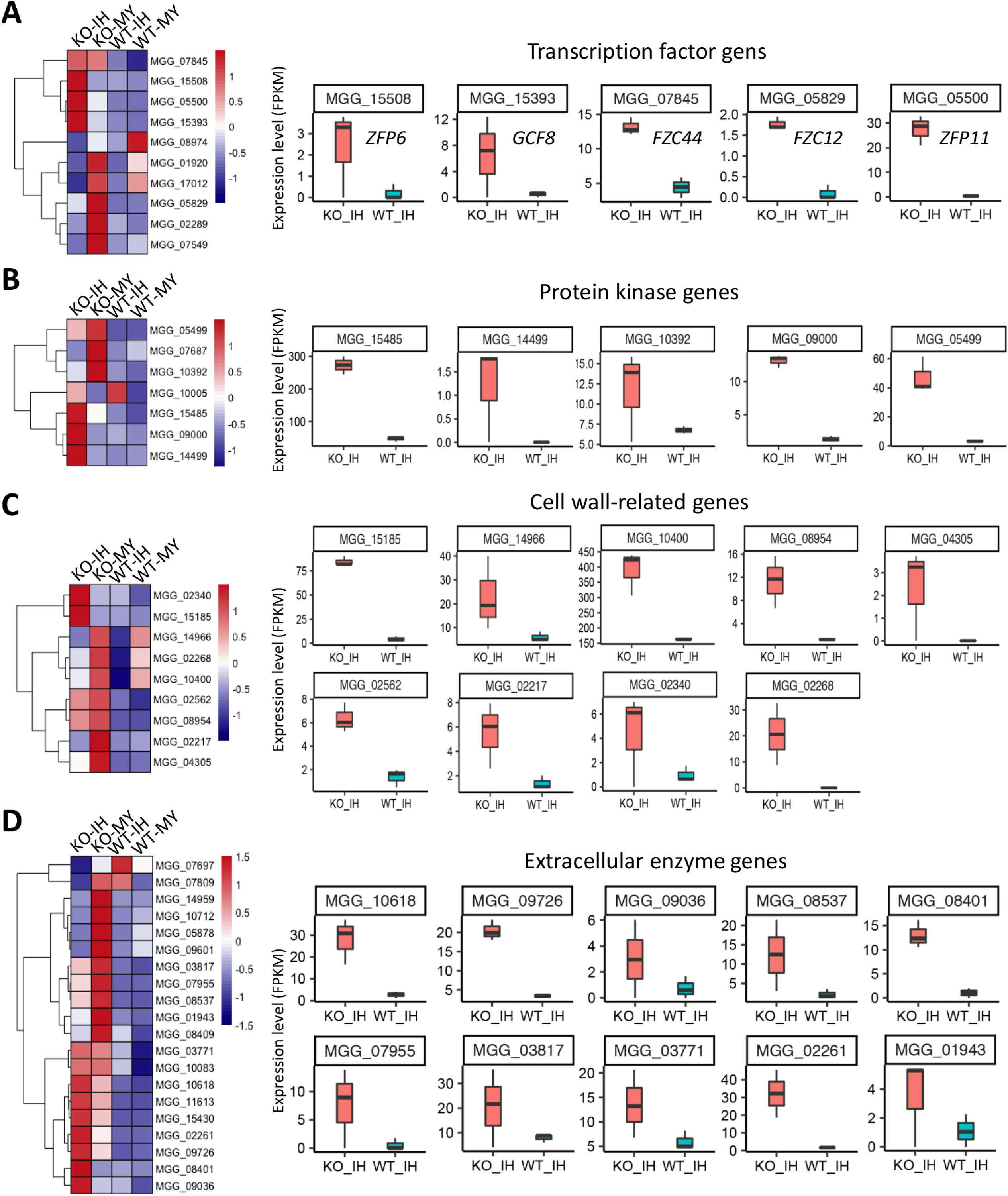
PRC2 represses secondary metabolites synthesis genes at biotrophic stage. (A) Heatmap shows expression of the metabolites synthesis genes in different samples. KO-IH, 24 hpi-invasive hyphae of the Δ*suz12* mutant; WT-IH: 24 hpi-invasive hyphae of WT; KO-MY, mycelium of the Δ*suz12* mutant; WT-MY: mycelium of WT. (B) Expression level of cytochrome P450 genes in KO-IH and WT-IH. (C) Expression level of polyketide synthase genes in KO-IH and WT-IH. (D) Expression level of other metabolites synthesis genes in KO-IH and WT-IH. (E) PRC2-repressed synthesis processes through regulating metabolites synthesis genes.

In total, at least 54 PRC2 target genes were annotated as secondary metabolite metabolism genes, including 25 cytochrome P450 genes, 8 polyketide synthase (PKS) genes, as well as 23 other genes. Seven cytochrome P450 genes encode isotrichodermin C-15 hydroxylase were regulated by PRC2 (Fig. 7A-D; Table S3). Isotrichodermin C-15 hydroxylase is involved in N, N-bisformyl dityrosine synthesis, which is related to deoxynivalenol (DON) synthesis in *Fusarium*, and important for necrotrophic growth. Five cytochrome P450 genes encoding lanosterol 14-alpha-demethylase ERG11 and two genes encoding C-22 sterol desaturase ERG5 were also enriched with H3K27me3 and mainly repressed by PRC2 (Fig. 7A and B; Table S3). The ergosterol biosynthesis pathway is required for generating ergosterol, a major component of the fungal plasma membrane (33). This ergosterol biosynthesis pathway is specific to fungi, and has been found to be an interesting target of antifungal drugs (34), especially the cytochrome P450 family Erg11 protein. Some polyketide synthase (PKS) genes were enriched with H3K27me3 and significantly decreased in expression at biotrophic growth stage. PKSs are a family of multi-domain enzymes that produce polyketides, which a large class of secondary metabolites (35). All these PRC2 target PKS genes were highly expressed in the Δ*suz12* mutant at biotrophic stage.

Many other secondary metabolites synthesis genes were also targeted and repressed by PRC2-mediated H3K27me3 (Fig. 7A, D and E). These include genes involved in biosynthesis of mycotoxin, alkaloid, steroid, lovastatin, gibberellin, ferricrocin, flavonoid, zeaxanthin, citrinin, mycophenolic acid, urocanic acid, pyoverdine/dityrosine, mycocerosic acid, paspaline, and monolignol, etc (Fig. 7E). Similarly, most of these genes were depressed in WT, but highly expressed in the Δ*suz12* mutant at biotrophic stage (Fig. 7A and D). All above data indicating that, PRC2-mediated H3K27me3 plays a key role in depressing secondary metabolites biosynthesis at biotrophic stage, which should be important for biotrophic growth during *M. oryzae* infection.

### PRC2 complex orchestrates effector-encoding genes expression during biotrophic growth stage

During infection, *M. oryzae* secreted many effector proteins to interact with host and facilitate its infection. How effector genes are elaborately regulated is largely unknown. Interestingly, we found many effector genes were enriched in the H3K27me3 targets at biotrophic growth stage. In total of 1701 H3K27me3 modified genes, 176 were effector-encoding genes (Table S4). Clustering analysis indicated that, in WT, most of these effector genes were repressed in mycelium, while around a half of them were significantly increased in expression in biotrophic stage (Fig. 8A). More interestingly, we noticed that another half of these effector genes which were evidently repressed in the biotrophic stage, but were significantly de-repressed in the Δ*suz12* mutant (Fig. 8A), indicating these genes were regulated by PRC2-mediated repression.

**FIG 8.**
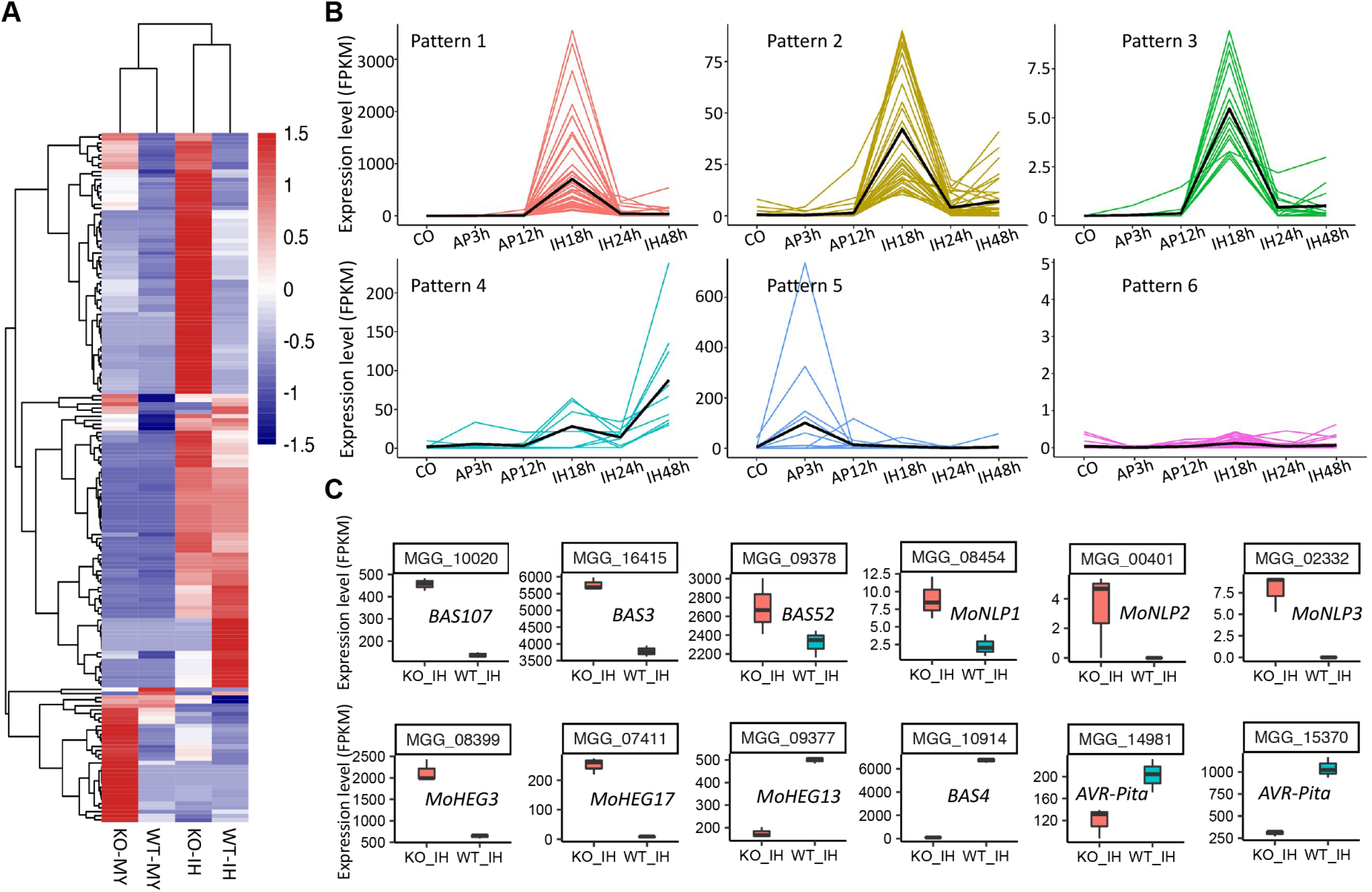
PRC2 regulates effector expression at biotrophic stage. (A) Heatmap shows expression of 176 H3K27me3 modified genes in different samples. KO-IH, 24 hpi-invasive hyphae of the Δ*suz12* mutant; WT-IH: 24 hpi-invasive hyphae of WT; KO-MY, mycelium of the Δ*suz12* mutant; WT-MY: mycelium of WT. (B) Expression of PRC2 complex genes at different developmental stages according to RNAseq data. MY, mycelium; CO, conidium; AP3H and AP12H, appressorium at 3 hpi and 12 hpi; IH18H, IH24H and IH42H, invasive hyphae at 18 hpi, 24 hpi and 42 hpi. (C) Expression level of known effector genes in KO-IH and WT-IH.

In order to characterize how these effectors are regulated during infection, we performed RNAseq analysis by using different samples collected from different developmental stages of the WT strain, including mycelium, conidium, immature appressorium (AP3h), mature appressorium (AP12h), penetration process/primary invasive hypha (IH18h), biotrophic invasive hypha (IH24h), and necrotrophic invasive hypha (IH48h) collected from barley leaves. To our surprise, we found expression of these effector genes can be clearly classified into several patterns (Fig. 8B). As shown in Fig. 9b, most effector genes expressed as pattern 1-3, in which their expression were repressed in mycelium, conidium and appressorium stage, but significantly increased at penetration stage when primary invasive hyphae initialized (IH18h). More interestingly, their expressions were sharply decreased at biotrophic stage (IH24h) and later stage, suggesting these genes function during penetration stage, but be repressed at biotrophic stage (Fig. 8B), which is modulated by PRC2-mediated repression. In pattern 4, several effector genes were also repressed at biotrophic stage, but again de-repressed at necrotrophic stage (IH48h), indicating these effectors play important roles in necrotrophic stage. In pattern 5, several effector genes were repressed in all tested stages except immature appressorium, suggesting they were required for this stage. In pattern 6, these effectors were repressed in all stages (Fig. 8B). Interestingly, above mentioned PKS gene *MGG_17768*, was also a known AVR gene, *ACE1* (confers AVR toward rice containing the R gene Pi33), which is a unique *AVR* gene. *ACE1* encodes a hybrid polyketide synthase-non-ribosomal peptide synthetase (PKS–NRPS) (36), probably function in synthesizing secondary metabolite recognized by the Pi33 gene product (37). It has been found that *ACE1* expression is tightly coupled to the appressorium-mediated penetration of the host cuticle (38), which is consistent with this study. Therefore, PRC2-mediated epigenetic control of the effector genes could be a universal mechanism help the fungi evade host immune recognition.

**FIG 9.**
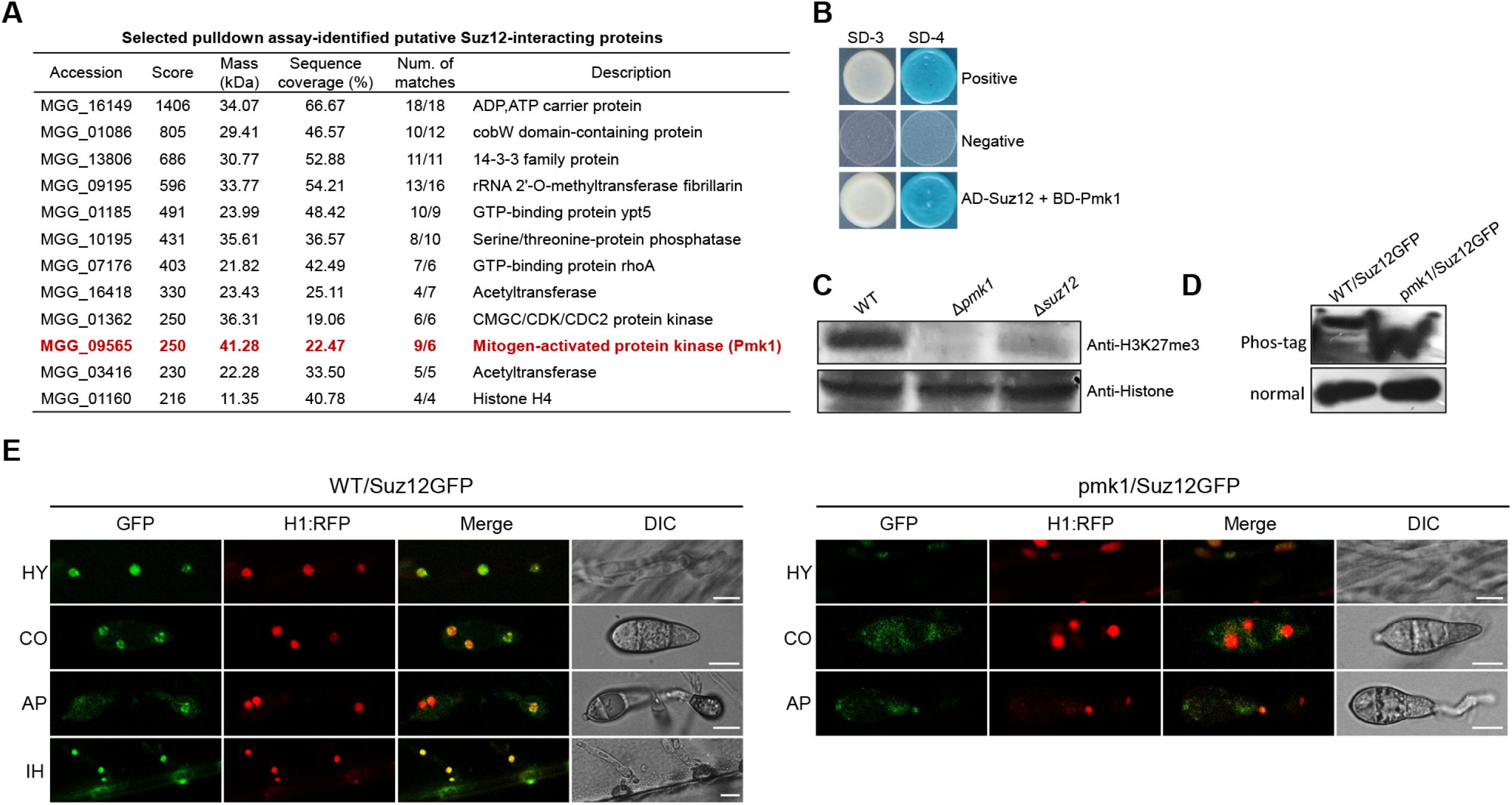
PRC2 is regulated by Pmk1. (A) List of pulldown assay identified Suz12-interacting proteins. Number of matches (Num. of matches) show two experiment replicates. (B) Yeast two-hybrid assay for the interaction between Suz12 and Pmk1. The interaction between pGBKT7-53 and pGADT7-T was taken as the positive control, while the interaction between pGBKT7-Lam and pGADT7-T was taken as negative control. BD, pGBKT7; AD, pGADT7. (C) Western blotting analysis showed histone H3K27me3 modifications in Δ*pmk1* mutant was disappeared. Total protein extracted from *M. oryzae* strains was subjected to 15% SDS polyacrylamide gel electrophoresis, and probed with antibodies against H3K27me3, and H3 histone. (D) Suz12 is phosphorylated by Pmk1. Suz12-GFP from wild-type and the Δ*pmk*1 mutant cell extracts were subjected to Phos-tag SDS-PAGE and normal SDS-PAGE followed by immunoblotting with anti-GFP. (E) Subcellular localization of Suz12-GFP in wild type and the Δ*pmk1* mutant in mycelium and conidium. WT/Suz12GFP, the wild-type strain expressing Suz12-GFP; pmk1/Suz12GFP, the Δ*pmk1* mutant strain expressing Suz12-GFP. Bars, 10 μm.

Some reported effectors genes such as *AVR-Pik* (39), *AVR-Pita* (40,41), *BAS107, BAS3, BAS4, BAS52* (24), *PWL3* (42), *MoNLP1, MoNLP2, MoNLP3* (43,44), *MoHEG3, MoHEG17, MoHEG13* (45) were included in these H3K27me3 targets. Among them, *BAS107, BAS3, BAS52, MoNLP1, MoNLP2, MoNLP3, MoHEG3, MoHEG17* were expressed as pattern 1-3, and significantly de-repressed in the Δ*suz12* mutant (Fig. 8C), suggesting they were regulated by PRC2-mediated repression at biotrophic stage. Avirulent gene AVR-Pik was expressed as pattern 6, which was low-expressed at all stages. However, there were also some effector expressions were increased in the Δ*suz12* mutant, including *MoHEG13*, *BAS4*, as well as two *AVR-Pita* genes (Fig. 8A and C). These suggested that PRC2 may also mediate gene activation, however, whose mechanism is unknown in the present.

To determine the correlation between abundance of PRC2 genes and effectors expression, we subsequently examined the transcription profile of PRC2 complex genes using quantitative real-time PCR (qRT-PCR). The results showed that, these genes seem low expressed in conidium and penetration stage (18 hpi) but highly expressed in appressorium formation and invasive growth stage (24 hpi), and subsequently low expressed again in necrotrophic stage (48 hpi) (Fig. 1E). This is consistent with previous result that PRC2 shows peak at 24 hpi, but low expression at 18hpi, while PRC2-supressed genes are highly express at 18hpi but decreased at 24hpi. Taken together, these results suggest correct expression of effectors are important to facilitate successful biotrophic growth and further colonization.

### Pmk1 regulates H3K27me3 through Suz12

It is now clear that PRC2 is fine tuned for infection of *M. oryzae*, but the up-stream regulatory mechanism is still unclear. Considering that Pmk1 plays key roles in biotrophic growth and cell-to-cell movement in *M. oryzae*, which is quite similar to PRC2, we speculate that PRC2 could be regulated by Pmk1. Interestingly, a pulldown assay by using Suz12-GFP as a bait, we identified Pmk1 was a putative Suz12-interacting protein (Fig. 9A). The interaction was subsequently confirmed by yeast two hybrid experiment (Fig. 9B). More interestingly, the H3K27me3 modification was totally disappeared in the Δ*pmk1* mutant (Fig. 9C), which is similar to that in the PRC2 genes’ deletion mutants, indicating that Pmk1 is also required for histone H3K27me3 modification. We further tested whether Pmk1 is involved in phosphorylation of Suz12. We analyzed the Suz12-GFP from the wild-type strain and Δpmk1 mutant on gels containing the Phos-tag, which retards the mobility of phosphoproteins. The reduced mobility form of Suz12-GFP was present in the wild-type strain but not in the Δ*pmk1* mutant (Fig 9D). This result suggested that Pmk1 could regulate Suz12 protein through phosphorylation. Further, we also observed whether Pmk1 can regulate subcellular localization of Suz12. In all tested samples, Suz12-GFP was clearly located in the nucleus, while evidently much weaker in the Δ*pmk1* mutant, and obviously tend to locate in the cytoplasm (Fig. 9E), suggesting Pmk1 regulate nuclear localization of Suz12. Because the Δ*pmk1* mutant can’t form invasive hyphae, we can’t detect subcellular localization of Suz12 in invasive hyphae. Together, Pmk1 can coordinate functions of PRC2 by regulating nuclear localization and protein abundance of Suz12, and therefore coordinate biotrophic growth and infection of *M. oryzae*.

## DISCUSSION

The roles of PRC2 complex have been identified and studied in several fungi, including *F. graminearum*, *F. fujikuroi*, *Z. tritici*, *C. neoformans N. crassa*, and *E. festucae* (16,17,18,19,20,21). In these studies, it has been shown that H3K27me3 correlates with gene silencing and plays a role in development, pathogenicity, and transcriptional regulation of secondary metabolite gene clusters (17,19). Recently, Functions of PRC2 was also revealed in *M. oryzae*, which indicated that PRC2-mediated H3K27me3 contributes to regulation of genes important during host infection, especially the effector proteins (26). However, it is far more unclear what’s the role of PRC2 complex during fungi-plant interactions. None of these studies used *in planta* fungal samples, instead by the mycelium samples, for testing, due to difficulty in collecting enough fungal samples. In this study, we collected the invasive hyphae samples from the *M. oryzae* infected barley epidermis cells at 24 hpi, which let us be able to enrich the fungus sample and successfully perform the ChIP-seq and RNAseq analyses. Therefore, we can elaborately determine regulatory mechanisms of PRC2-mediated H3K27me3 during biotrophic growth of *M. oryzae*. Using these high-quality ChIP-seq and RNAseq data, as well as biological analyses to the PRC2 genes’ deletion mutants, we were able to reveal that PRC2 is essential for *M. oryzae* biotrophic growth though reprogramming gene expression pattern, and uncover the regulatory mechanisms. According to our study, change of gene expression between WT and Δ*suz12* in mycelia is not too much, but significantly different is found in biotrophic invasive hyphae, indicating more important roles of PRC2 in biotrophic stage.

Biological analysis demonstrated that PRC2 is essential for biotrophic growth of *M. oryzae*, but how PRC2 can regulate biotrophic growth? ChIP-seq and RNAseq data analysis showed at least several possible regulatory mechanisms of PRC2-mediated biotrophic growth. First, PRC2 can repress expression of genes encoding cell wall-related proteins, CWDEs, secondary metabolites biosynthesis proteins, as well as effector proteins. These suggested that in order to facilitate biotrophic growth, *M. oryzae* need to suppress unnecessary gene’s expression. During penetration process, *M. oryzae* needs to keep cell wall rigidity to help penetration. At the same time, it also secreted kinds of CWDEs such as celluloses, cutinases, metalloproteases, exoglucanases, and xylanases etc., to digest plant cell wall and facilitate penetration (32). After penetration, the fungus needs to quickly establish biotrophic growth, and switches gene expression patterns to save energy. We propose that PRC2-mediated epigenetics regulation plays key roles at this time. Consistent with this view, many PRC2 bound cell-wall synthesis genes and extracellular enzyme genes are regulated by PRC2-mediated repression at biotrophic growth stage. Second, the fungus also needs to suppress secondary metabolites synthesis which mainly occur in necrotrophic growth stage. Consistently, many cytochrome P450 genes, PKS genes as well as many other secondary metabolites synthesis genes were targeted and suppressed by PRC2. Some transcription factors and protein kinases may be also involved in regulating necrotrophic growth process and were repressed by PRC2 at biotrophic stage. Third, PRC2 also suppresses many effector genes, which is much more important for evading host recognition during interaction. We found many effector proteins were directly repressed by PRC2 at biotrophic growth stage, some of which were sharply reduced expression from a very high expression level. It seems that when these effectors finish their roles in suppressing host immune system, they should be quickly repressed to evade host recognition. On the other hand, effector production may be elaborately regulated to avoid wasting energy under conditions when the gene products are not needed. We show that even between biotrophic growth initiation and establish, some effectors were also exactly regulated. PRC2-mediated epigenetic repression therefore function as a key switch to finely tune effector gene expression for biotrophic growth.

However, another remain question is that the upstream mechanisms regulating PRC2 in fungi is largely unknown previously. Here we show that the protein kinase Pmk1 may play a key role in fine-tuning PRC2-mediated H3K27me3 for genome-wide gene remodeling (Fig. 10). It is known that Pmk1 plays central roles in regulating biotrophic growth during *M. oryzae* infection, especially in expression of secretory fungal effector proteins, inhibiting host immune defense (Wilson and Talbot, 2009; Sakulkoo et al., 2018), but its downstream mechanism is still unclear. Previous hypothesis is that it can regulate central transcription factor such as Mst12 for gene remodeling (Wilson and Talbot, 2009; Park et al., 2002). Now in this study, we provided another possibility that, Pmk1 can also regulate epigenetic switch such as PRC2-mediated H3K27me3 for gene expression remodeling. It is interesting to test if any other epigenetic target of Pmk1 in the future.

**FIG 10.**
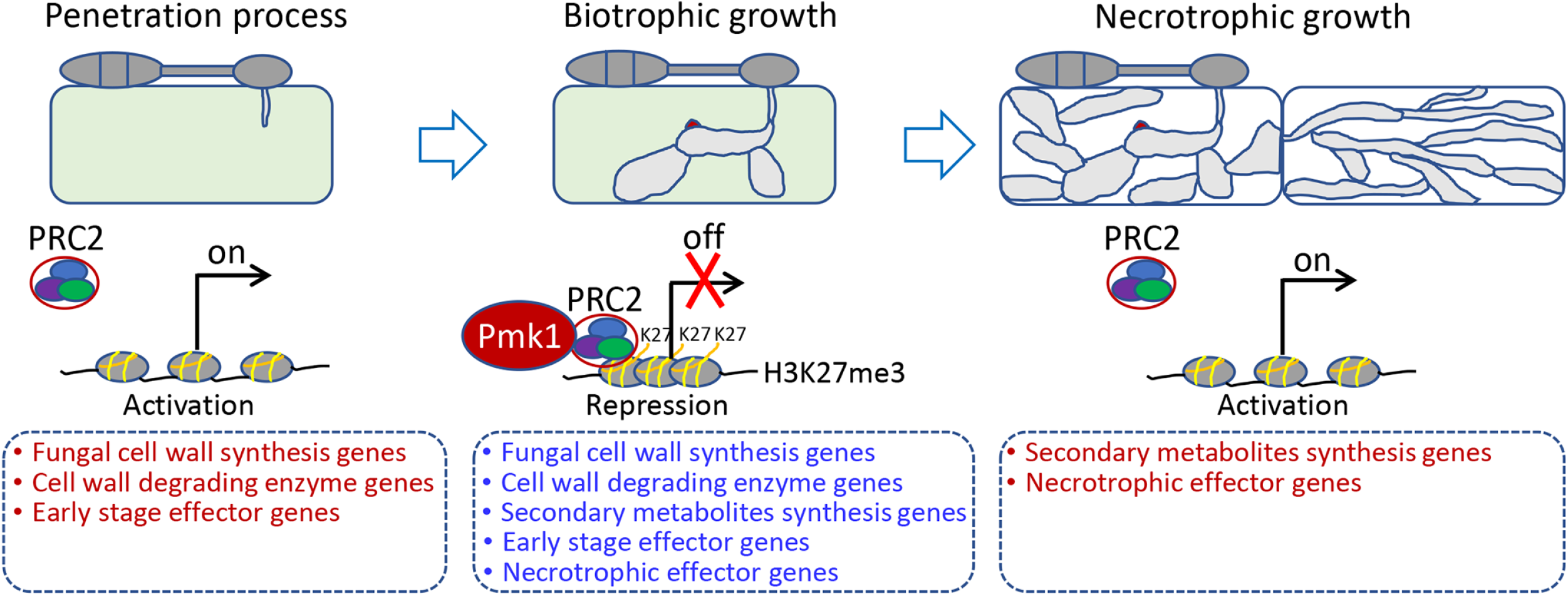
A regulatory model of PRC2-mediated repression in regulating of biotrophic growth in *M. oryzae*.

Taken together, we propose a model that how PRC2 coordinates biotrophic growth through repressing different gene expressions (Fig. 9). PRC2 ultimately functions to repress occupied genes in biotrophic growth stage and that these genes may be especially toxic, wasteful, or host sensitive for transcriptional activation during biotrophic growth. PRC2 binding during biotrophic growth represses key developmental regulators that are expressed and required for other stages. Obviously, genes involving in fungal cell wall synthesis, extracellular CWDEs, and early-stage effector genes at penetration processes are quickly repressed by PRC2 at biotrophic stage. Before switching into necrotrophic growth, genes related to secondary metabolites synthesis, and necrotrophic effector genes are also required to be repressed by PRC2.

Fungal strains containing sequence variations or changed expression patterns of AVR genes can help them evade plant R protein recognition, resulting in a failure of plant immunity (46). Epigenetic code could provide more rapid way to respond to environmental stress than DNA sequences themselves for fungal evolution. Some AVR effectors, including ACE1, AVR-Pita, and AVR-Pik, were identified as PRC2 targets. A recent study also found that PRC2-mediated H3K27me3 is required to maintain the naturally silenced state of Avr1b in a *Phytophthora sojae* strain to evade host recognition (47). How fungal pathogens adapt to host plants through epigenetic regulation remains largely unexplored. It is interesting to further reveal the regulatory mechanisms of PRC2 on AVR genes in the future, by using different field strains and rice cultivars for testing.

## MATERIALS AND METHODS

### Strains and culture conditions

All the wild-type strain of *M. oryzae* used in this study is P131 (48). The fungal strains were grown on Oatmeal Tomato Agar (OTA) plates at 28°C. We incubated the mycelia in CM liquid culture at 28°C for 36 h for DNA extraction and protoplast isolation. Colony growth and conidiation were operated as described according to Chen et al 2014. To examine the virulence and determine the infection process, conidia were produced and collected from 7-day-old OTA cultures. For stress sensitivity test, strains were inoculated on CM plates supplemented with different stress agents (0.2 mg/ml Congo Red [CR], 0.1 mg/ml Calcofluor White [CFW], 0.005% Sodium dodecyl sulfate [SDS], 0.5 M NaCl, and 10 mM H_2_O_2_), and the colony diameters were measured at 120 hours post-inoculation (hpi). To observe cell lengths of the hyphal tips, mycelia in CM liquid culture were stained with 10 μg ml^−1^ CFW (Sigma-Aldrich, USA) for 10 min in the dark, and observed under a fluorescence microscope (Nikon Ni90 microscope, Japan).

### Virulence test and infection process observation

Four-week-old rice seedlings (*O. sativa* cv. LTH) and 1-week-old barley leaves (*Hordeum vulgare* cv. E9) were used for virulence test. Conidial suspensions of different strains (5 × 10^4^ conidia/mL in 0.025% Tween 20) were sprayed onto plant leaves, which were subsequently incubated in full humidity condition at 28°C. Five days later, the disease symptoms were observed and photographed. To observe infection process, conidial suspension (1 × 10^5^ conidia/mL) were dropped onto barley leaves, which were subsequently incubated in full humidity condition at 28°C. The lower barley epidermis was used for observing appressorium-mediated penetration at 18 hpi, biotrophic growth at 24 hpi, and necrotrophic invasive growth at 48 hpi under a microscope. ROS accumulation in host cells infected by different strains at 36 hpi was observed by staining the cells with DAB (Sigma-Aldrich) as described (48).

### Western blot analysis

To detect H3K27 methylation levels in different strains, fungal mycelia samples were used to extract total proteins. Western blotting was performed as previously described (49), using antibodies of anti-H3K27me1, anti-H3K27me2 and anti-H3K27me3 (Jingjie, Hangzhou, China), and anti-H3 (Jingjie, Hangzhou, China) was used as control.

### ChIP and data analysis

ChIP-seq experiment was performed as previously described with modifications (50,51). Invasive hyphae of WT and Δ*suz12* were harvested by collecting fungus infected barley epidermis at 24 hpi. The samples were cross-linked with 1% formaldehyde in CM liquid culture and shaking for 30 minutes at room temperature. Crosslinking was stopped by adding 125 mM glycine. Chromatin was sheared by sonication for 5 times to obtain DNA approximately 100-500 bp fragments. 50 μl of protein A (Dynabeads) (Invitrogen, Carlsbad, CA) was incubated with 5 ml antibody H3K27me3 in 1 ml of ChIP dilution buffer at 4 °C for 5 h. Control samples were parallelly handled but without adding the antibody. Protein-beads were collected by a magnet and washed for four times with ChIP dilution buffer, and then incubated overnight with 1 ml of chromatin dilution buffer at 4 °C. The antibody–chromatin complex was washed, eluted and de– cross-linked. DNA was recovered by phenol–chloroform extraction and ethanol precipitation in the presence of glycogen and resuspended in 20 ll of distilled water. The samples were used to generate the sequencing library with Illumina HiSeqTM2000 for 50nt pair-end sequencing.

The IP and control raw reads were checked by fastqc (version 0.11.5). Low quality reads were filtered using Trimmomatic (version 0.36). The clean reads from each library were aligned against to the *M. oryzae* reference genome version 8 using BWA (version 0.7.15). PCA and Spearman Correlation analysis was performed to determine similarity and distance between replicates. MACS2 software (version 2.1.1.20160309) was used to define peak by default parameters (bandwidth, 300bp; model fold, 5, 50, qvalue < 0.05). Wig files produced were used for visualization in IGV (version 2.8.4). deeptools (version 2.5.4) was used to determine distributions of reads to 2 kb distance to TSS (transcriptional start sites). To define the association between genes and H3K27me3 modification, summit position of each peak was used, and the closest TSS of corresponding gene was collected. Raw data of the ChIP-seq and RNA-seq were submitted to Sequence Read Archive (SRA) database with a combined accession number PRJNA721132.

### RNA sequencing and data analysis

Invasive hyphae of WT and Δ*suz12* were harvested by collecting fungus infected barley epidermis at 24 hpi. Two biological replicates for each strain were performed. Total RNA was extracted using TRIzol (Invitrogen, Karlsruhe, Germany) according to manufacturer’s instructions. The extracted RNA was treated with DNase and cleaned up using the RNA Clean & Concentrator-25 Kit (Zymo Research, Freiburg, Germany). RNA samples were then used for sequencing on an Illumina Hiseq2000 platform. Low quality reads were filtered by Trimmomatic (version 0.36), and checked by fastqc (version 0.11.5). Filtered reads were aligned to the *M. oryzae* reference genome version 8 using subread, and defined reads counts using featureCounts. FPKM values were generated after normalization. PCA and Pearson analysis were performed to determine distance between three replicates. edgeR package was used for differential expressed genes analysis, using cut-off as FDR < 0.05, and fold change > 2. Phylogenetic analysis was generated and visualized using mega7. To predict effectors, SignalP and EffectorP were used by inputting peptides sequences of *M. oryzae*.

## ACKNOWLEDGMENTS

This work was supported by the National Natural Science Foundation of China (Grants 31871909 and 32072365 to X.L.C.) and the Open Research Fund of the State Key Laboratory of Hybrid Rice (Hunan Hybrid Rice Research Center) (2019KF04 to X.L.C.).

X.L.C. conceived the project, X.C., A.H., C.L., and Z.R. performed the biological experiments, B.T. performed the bioinformatic analyses, J.X., L.Z., H.L., J.H. interpreted results, X.L.C., and B.T. wrote the paper. All authors edited and approved the paper.

## Supporting Information

**FIG S1** Phylogenetic trees of three PRC2 proteins. Phylogenetic trees were constructed by the neighbor-joining (NJ) method and the Kimura 2-Parameter model with bootstrap resampling by the MEGA 6 package.

**FIG S2** Protein domains of three PRC2 proteins.

**FIG S3 Gene deletion of PRC2 complex genes.** (A) Gene replacement of PRC2 genes through a split-marker approach. White bars represent genomic regions upstream and downstream of the coding sequence that were amplified and fused to segments of the hygromycin phosphotransferase (HYG) cassette. (B) PCR verification of the flanking sequences beside the replacement fragment by using primer pairs of LCK/HCK-up and RCK/HCK-down. (C) RT-PCR verification by amplifying the expression of the gene in the transformants and the wild-type strain. For (B) and (C), KO1-1 and KO1-2: *EED1* deletion mutants; KO2-1 and KO2-2: *EZH2* deletion mutants; KO3-1 and KO3-2: *SUZ12* deletion mutants.

**FIG S4 PRC2 complex is involved in growth and conidiation of *M. oryzae*.** (A) Colony morphology of the wild type P131 and deletion mutants of the PRC2 complex genes. All strains were observed on oatmeal tomato agar (OTA) plates at 28°C for 5 days. (B) The colony diameters were measured and subjected to statistical analysis. Error bars represent standard deviation and asterisk represents significant difference (P < 0.01). (C) Cell length of the hyphae tips in the wild type P131 and deletion mutants of the PRC2 complex genes. The triangles indicate septa between cells. (D) Statistical analysis of cell length. Error bars represent the standard deviation and asterisks represent significant difference among strains (P < 0.01). (E) Conidial and conidiophore formation were observed under a light microscope. The indicated strains were grown on OTA plates for 5 days. (F) Statistical analysis of conidiation. Error bars represent the standard deviation and asterisks represent significant difference among strains (P < 0.01).

**FIG S5** Co-relation of ChIPseq result in different samples. (A) Co-relation analysis to ChIP-seq data of different samples. (B) Principal component analysis (PCA) to ChIP-seq data of different samples.

**FIG S6** Co-relation of RNAseq result in different samples. (A) Co-relation analysis to RNAseq data of different samples. (B) Principal component analysis (PCA) to RNAseq data of different samples.

**TABLE S1 Information of the H3K27Me3 ChIP-seq peak reads.**

**TABLE S2 Expression of RNAseq identified genes.**

**TABLE S3 List of PRC2-targeted important genes.**

**TABLE S4 Expression of 176 PRC2 regulated effector genes at different developmental stages.**

